# Linking reaction time variability to physiological markers of arousal across timescales

**DOI:** 10.64898/2026.03.19.713034

**Authors:** Deepa Issar, Emily E. Skog, Madison R. Grigg, Jana M. Kainerstorfer, Matthew A. Smith

## Abstract

Reaction time is a measure of the speed of our response to stimuli in the environment. Even for a well-trained task, a subject’s reaction time varies. One source of this variability is internal state fluctuations (such as changes in arousal). There are few studies that systematically quantify the extent to which reaction time varies across different timescales and link this to measures of systemic physiology associated with arousal. In much of the literature, it is assumed but not demonstrated that behavioral and systemic measurements associated with arousal will be consistently linked because both estimate a common underlying arousal process. In this work, we examined this assumption by simultaneously measuring reaction time, heart rate, and pupil diameter in rhesus macaque monkeys performing several visual tasks over hours and across hundreds of sessions. We found a portion of the variability in reaction time could be linked to systemic physiological signatures of arousal on fast timescales from second to second and slower timescales from minute to minute. This link between reaction time and systemic physiology was also present for different biomarkers of arousal (heart rate and pupil). However, the strength of this relationship varied depending on the arousal biomarker. Our findings support the conclusion that there are multiple arousal mechanisms that act simultaneously to influence behavior and multiple timescales at which they operate.

## INTRODUCTION

If a rattlesnake slithers across the trail during a hike, reacting quickly to avoid stepping on it is critical. Reaction time (RT) measures the duration between a prompting stimulus and the onset of the resulting action. In laboratory settings, it is typically defined as the time between a task-specific go cue and the initiation of a movement, such as a saccade or reach. The go cue must be transformed into a neural signal that traverses sensory, cognitive, and motor brain areas before being transmitted to muscles to become a motor response. As a result, numerous brain and body processes can have an impact on RT.

RT is typically split into non-cognitive and cognitive components (1–3). The non-cognitive components include processes outside of decision-making that affect RT. This includes sensory transmission time, which is the time for a stimulus in the environment to be transformed into neural signals in the sensory system. For a visual stimulus this lag can be from 30 to 60 ms to activate neurons in visual cortex (4,5). Non-cognitive components of RT also include motor transmission time, during which neural commands in motor cortical areas propagate to muscles, and muscle activation time, the duration from muscle activation to visible muscle contraction. For a saccade, a rapid eye movement to a target that relies on the coordination of several eye muscles, the combined motor transmission and muscle activation time can be around 20 to 60 ms (6,7). For skeletomotor actions, such as forelimb movements, this motor to muscle period can be a hundred or more milliseconds (8,9).

The cognitive component of RT consists of processes that influence decision time, the epoch during which the subject interprets incoming sensory information and resolves to act on that information. Decision time is often modeled as a period of noisy evidence accumulation that varies based on the rate of accumulation, the threshold at which a decision will be made, and other parameters (1,2,10–12). This period can be hundreds of milliseconds and depends greatly on the task type (13). In tasks with a go cue that is ambiguous or close to the subject’s perceptual threshold, decision time is longer than in simple detection tasks with a clear go cue (14). Alternatively, decision time can also be longer because the subject has a conservative threshold for making the decision (15).

RT can vary considerably on matched repeated trials even for a well-trained subject on a simple task (13,16,17). Numerous psychological and environmental factors contribute to this variability in RT. Increased attention, which is the dedication of cognitive resources to a particular spatial region or object, results in faster RTs with less variability (18–20). Motivational factors, such as reward and urgency, also influence RT, with larger rewards leading to more vigorous movements and faster RTs (21–23) or more cautious movements and slower RTs (24,25). Increasing urgency by encouraging subjects to respond as quickly as possible decreases RT (26–28). Vigilance and overall level of engagement can decrease over sustained tasks, leading to longer RTs (29–31). In a similar vein, slower RTs are associated with physical or mental fatigue (32,33). Alertness and fatigue are also tied to the 24-hour sleep-wake cycle known as the circadian cycle, and RT has been shown to vary with environmental factors like time of day (34,35).

Many of the phenomena that give rise to variability in RT, such as engagement, fatigue and alertness, can be captured under the umbrella term arousal. Broadly, arousal level reflects a state of internal readiness to engage with our environment (36,37) that fluctuates over seconds (38), minutes (39), and throughout the day (40). At the neural level, the waxing and waning of arousal appears to arise in part via the coordination of several brainstem nuclei, including the locus coeruleus (38,41,42). Measurement of activity in these brainstem nuclei is challenging, and their effects on the brain are diffuse and difficult to understand from observations of a small number of neurons in a single brain area.

Since arousal level is difficult to estimate directly from neural activity, behavior and physiology are frequently used as arousal proxies. Given the definition of arousal is connected to our readiness to respond, faster RTs are associated with elevated arousal. Some studies will solely rely on behavioral changes in RT as an indicator of fluctuations in internal state like arousal level (43–45), even though there are many influences on RT (described above). Alternatively, since arousal-linked brainstem nuclei are closely tied to systemic physiology, changes in pupil diameter (46–48) and heart rate (HR) (49–51) are also popular biomarkers of arousal. Similar to RT, these physiological measurements are influenced by more than arousal level (52). It is possible that shared fluctuations in a group of behavioral and physiological markers may constitute a more robust arousal signature. A few studies in the literature have begun to address this idea by quantifying the extent to which changes in RT can be explained by single measures of systemic physiology, such as heart rate (53,54), heart rate variability (55), pupil (56,57) and galvanic skin response (58). However, it is rare for any one study to measure multiple types of systemic physiology as well as capture them across multiple timescales in the context of carefully controlled psychophysical tasks. Without simultaneous physiology measurements across different timescales, we cannot extrapolate whether these arousal-linked behavior and physiology relationships generalize to different types of physiology and temporal scales.

Our goal was to understand the extent to which variability in saccadic RT over seconds, minutes, and days could be linked to systemic physiological measurements. We measured HR in rhesus macaque monkeys performing visual tasks. We found that some of the variability in RT from second to second and from minute to minute could be linked to changes in HR on the same timescale. Furthermore, we assessed how closely HR reflected changes in pupil diameter. We found these physiological measures were linked but could not always explain variability in RT to the same degree. Our work characterizes the relationship among various arousal-linked measurements and helps us interpret the connection between studies on arousal that use different arousal biomarkers.

## MATERIALS AND METHODS

We measured RT, HR, and pupil diameter from two adult male rhesus macaques (*Macaca mulatta*): subjects RA and AB. Prior to these experiments, animals were implanted with a titanium headpost using sterile surgical techniques under isoflurane anesthesia. Animals performed two visual tasks, a simple saccade task and a color change detection task (Fig 1). Subjects were rewarded with drops of water for looking to the correct target location. All subjects were well-trained on these tasks, which we assessed by stable task performance across sessions. Only correct trials were used for analyses. All experimental procedures were approved by the Institutional Animal Care and Use Committee of Carnegie Mellon University and were performed in accordance with the National Institutes of Health *Guide for the Care and Use of Laboratory Animals*.

**Fig 1.**
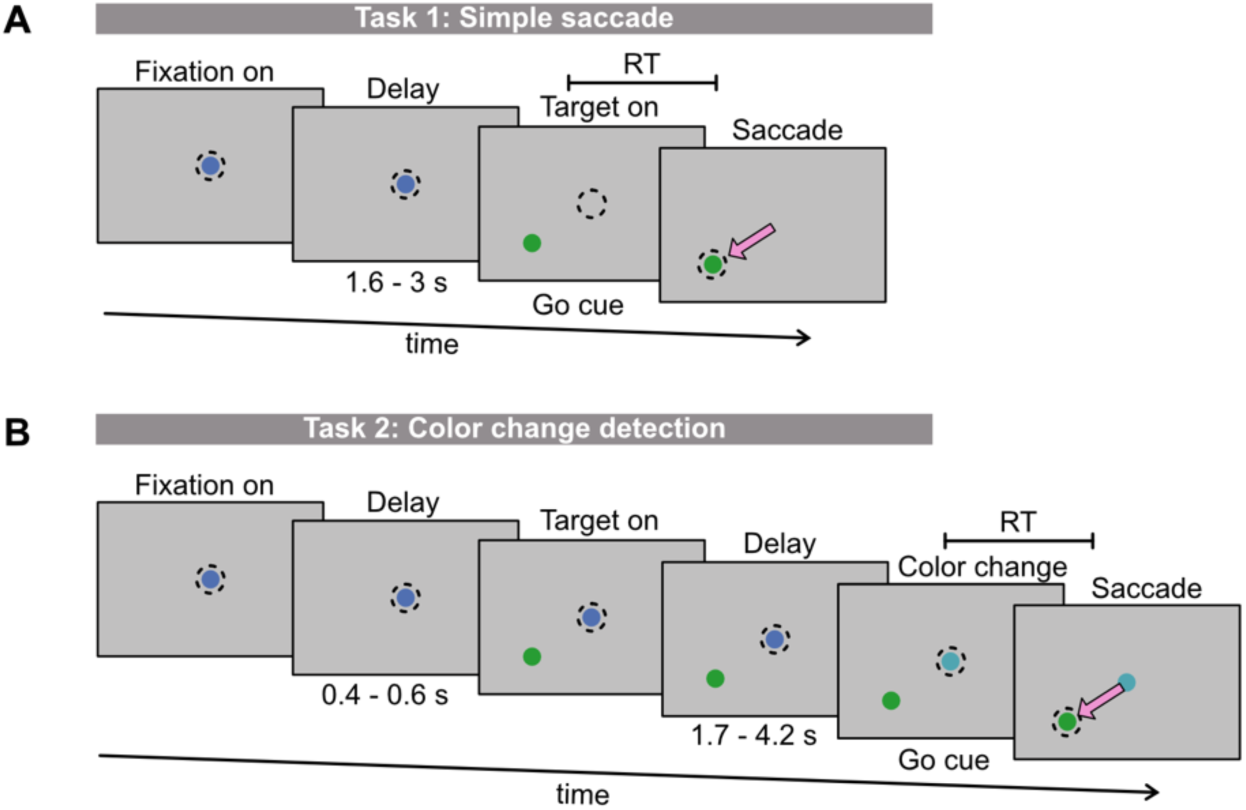
Reaction time (RT) was estimated in two types of visual tasks. Subjects fixated on a central dot (blue) for a random delay until the go cue appeared. RT was the time from the go cue until the subjects initiated a saccade. Subjects were rewarded for making quick and accurate saccades to the target (green). A) In task 1, the go cue was the simultaneous disappearance of the central fixation dot and appearance of the target. B) In task 2, the target appeared prior to the go cue which was when the central fixation dot changed color.

### Simple saccade (Task 1)

Subjects fixated on a dot at the center of the screen for a delay period (ranging from 1600 – 3000 ms across sessions). This delay was comprised by an initial random variable period (chosen uniformly from 400 to 600 ms), a fixed period that was a constant value for each session (ranging from 1200 to 2100 ms across sessions), and another random variable period (chosen uniformly from 0 to 300 ms). On a fraction of trials, during the fixed period of the delay a large checkerboard stimulus flashed on the screen; however, this did not provide the subject any information about the trial and was not relevant for the analyses in this work. After the delay period, the central fixation dot disappeared, and a target appeared at one of eight locations (set equally from fixation at angles spaced by 45°). This was the subject’s go cue to look at the target (Fig 1A). The subject failed the trial if they broke fixation prior to the go cue or did not quickly saccade to the target (<500 ms to initiate the saccade, <200 ms to complete the saccade) and accurately (within 2.2° of the target). Considering only failed trials in which the subject did not make an accurate saccade to the target after the go cue appeared (i.e. excluding broken fixation trials), subject RA’s average performance was 98.7 ± 1.8% across 72 sessions and on average he performed 1837 ± 575 correct trials per session. Subject AB’s average performance was 90.5 ± 4.4% across 54 sessions and on average he performed 1411 ± 267 correct trials per session.

### Color change detection (Task 2)

Subjects fixated on a dot at the center of the screen for a random variable delay chosen uniformly from 400 to 600 ms. Next, a target appeared at one of eight eccentricities (spaced in steps of 45°); however, unlike the simple saccade task, subjects had to continue to fixate for an additional delay period. This second delay was chosen from a random exponential distribution with a mean of approximately 2000 ms, a minimum value of 1700 ms, and a maximum value of 4200 ms. On a fraction of trials, during the first 1200 ms of the second delay a stimulus flashed on the screen; however, this did not provide the subject any information about the trial and was not relevant for the analyses in this work. After the second delay period, the fixation dot at the center of the screen changed color. This was the subject’s go cue to look at the target. The color change was a rotation in HSV color space (by 1°, 4°, 8°, or 10°), where a 360° rotation would return to the original color of the fixation dot (base color in HSV space: 240°). For rotations less than 180°, a large rotation in color space (e.g. 8°) was easier to detect than a small rotation (e.g. 1°). The difficulty varied from trial to trial. The subject failed the trial if they broke fixation prior to the go cue, looked at the target before the go cue (false alarm), did not look toward the target accurately (within 2.4° of the target) and quickly after the go cue (<500 ms to initiate the saccade, <200 ms to complete the saccade). Including all trial outcomes except those in which the subject broke fixation, Subject RA’s average performance was 64.5 ± 9.0% correct across 27 sessions and on average he performed 1354 ± 389 correct trials per session. Subject AB’s average performance was 70.3 +/− 5.3% correct across 20 sessions and on average he performed 796 ± 155 correct trials per session.

### Physiology measurements

#### Heart rate

We estimated HR from the pulsatile vascular signal measured using a near-infrared spectroscopy probe placed on the subject’s scalp (PortaLite; Artinis Medical Systems). The probe uses LEDs to shine light at 750 and 860 nm onto the scalp. By detecting the returning light and using the Modified Beer Lambert Law, we estimated changes in oxyhemoglobin over time (59). By filtering within a range of 1 to 3 Hz using a 2^nd^ order Butterworth filter and peak detection, we isolated peaks at the HR frequency.

#### Pupil diameter and eye tracking

We recorded eye position and pupil diameter monocularly using an infrared eye tracking system at a rate of 1000 Hz (EyeLink 1000 SR research). Saccadic RT was estimated as the time from the appearance of the go cue to the time when the subject’s eye left the fixation window (1.5° radius around the central fixation dot).

#### Estimating behavior and physiology on different timescales

While we can simply estimate the variability in the raw signals and the relationship between them over the session, we also wanted to assess these parameters on fast and slow timescales. We explored three different timescales.

#### Second to second timescale

To capture trends on the order of seconds and remove the influence of slow fluctuations in the mean, we took the value of the signal (e.g. HR, pupil, or RT) on each trial and subtracted the 100-trial moving average. This gave us a residual value for each trial over the session that captured changes in the signal occurring at timescales faster than 100-trials. For the simple saccade task (task 1), it took on average 8.5 ± 2.4 minutes to complete 100 correct trials. For the color change detection task (task 2), it took on average 13.3 ± 2.2 minutes to complete 100 correct trials. We refer to this timescale as ‘second to second’ because it includes fluctuations occurring from trial to trial over several seconds.

#### Minute to minute timescale

We estimated fluctuations on the order of minutes using the 100-trial moving average. We chose to use 100-trial bin sizes for the moving average because they were similar in size to the bins used in our previous work (39) to identify slow behavioral fluctuations over the session and we found similar results when testing other bin sizes ranging from 20 to 400 trials. The moving average was computed with non-overlapping bins. This gave us a mean value for each 100-trial bin over the session, where the number of bins was the total number of correct trials divided by 100. For the simple saccade task (task 1), there were on average 16 ± 5 bins per session and the sessions were on average 139.3 ± 47.9 minutes. For the color change detection task (task 2), there were on average 11 ± 4 bins per session and the sessions were on average 148.7 ± 44.9 minutes.

#### Day to day timescale

To estimate trends occurring on timescales of several hours (ultradian) and across days (infradian) that affected our session to session measurements, we computed a single mean value for each session. Since each session could be a different length and the beginning of the session tended to be quite variable, we used trials 200 to 300 of each session to compute the mean. This ensured that the mean best represented a similar time in each session without being influenced by things occurring prior to the start of the experiment. We tested other trial ranges from the beginning of the session and found the exact range chosen had minimal impact.

#### Statistical tests

For correlations, we evaluated statistical significance (i.e. a non-zero correlation) using a two-tailed t-test. For Pearson’s correlation values, we used the Fisher r-to-z transformation prior to significance testing across all sessions for a particular subject and task. We used partial correlation to assess the relationship between two of our signals when controlling for variance explained by the third. Lastly, to assess whether variability in RT and HR was influenced by time of day we used a binomial exact test. The goal was to assess whether separating sessions into one of two categories based on the time at which they began (an AM subset and a PM subset) decreased variability across sessions within each subset. This would indicate that some of the session to session variability was linked to time of day. We took the ratio of the standard deviation across the session subset to the standard deviation across all sessions, resulting in four values (two subjects with an AM and PM value for each subject). Using these values, we could test whether the proportion of values where this ratio was less than 1 (indicating standard deviation decreased when accounting for time of day) was greater than expected by chance.

## RESULTS

We recorded saccadic RT (correct trials only) while well-trained subjects performed a visual task (see *Methods*). Even though the animals could perform the task well in each session, we observed variability in RTs over time (Fig 2A, B). We aimed to both quantify this variability as well as assess how much of the variability could be linked to arousal level, which we estimated using HR (Fig 2C). The RT and HR on each correct trial over the session were negatively correlated in many sessions (r = −0.14, p <0.0001 for traces in Fig 2A and C). To examine whether this relationship was primarily driven by slow drifts in the measurements over minutes, faster fluctuations over seconds, or both, we estimated the relationship between our measurements on different timescales.

**Fig 2.**
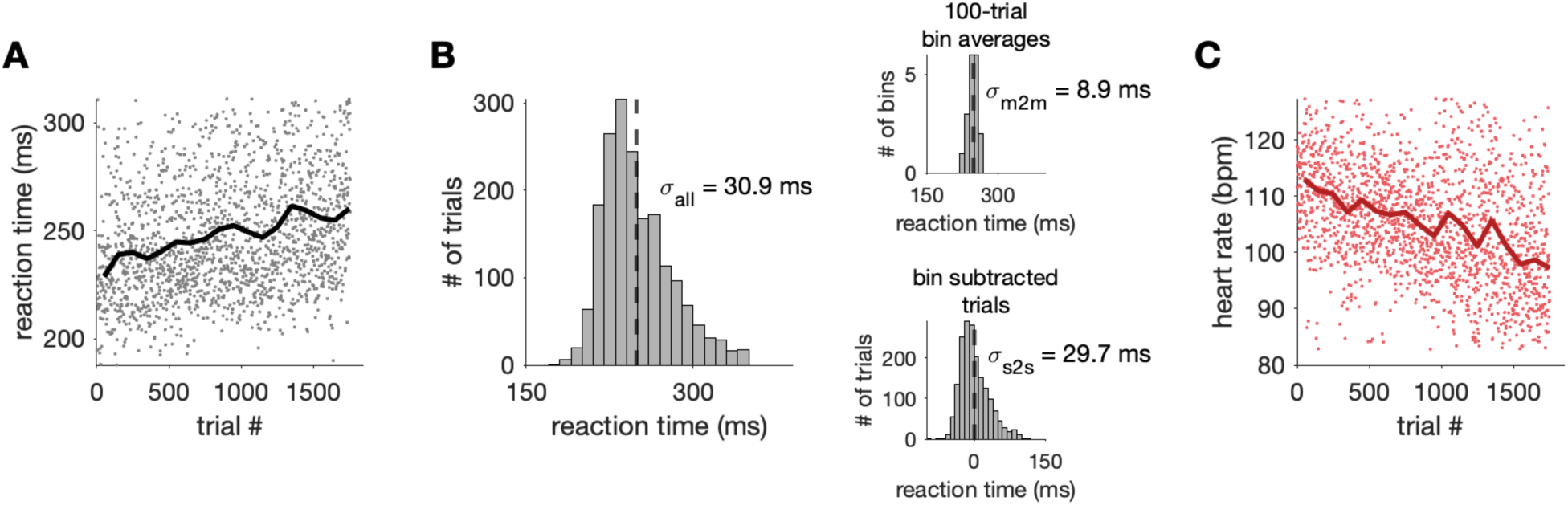
RT and heart rate (HR) varied over the session. One example session of the change detection task for Subject RA. A) RTs for all correct trials over time (black line: moving average using 100-trial bins). B) Distribution of correct trial RTs (dashed line: mean). Top inset: distribution of binned 100-trial average RTs (m2m: minute to minute timescale). Bottom inset: distribution of RT residuals after subtracting binned 100-trial average from all trials in that bin (s2s: second to second timescale). C) Same as in A) but for HR on each correct trial (red line: moving average).

We focused on three timescales: second to second, minute to minute, and day to day. Second to second variation was assessed from trial to trial in task performance, since each trial lasted 1.5 to 5 seconds and contained one behavioral response (saccadic RT). Minute to minute variation was measured over the course of large groups of adjacent correct trials (approximately 10 minutes) within individual sessions. Day to day variation was assessed between different sessions and therefore included both ultradian (< 24 hours) and infradian (> 24 hours) sources of variability.

We separated second to second and minute to minute fluctuations in the data by taking the moving average in 100-trial non-overlapping bins. The amount of time elapsed over this bin size varied depending on the task (simple saccade task: ∼8 minutes, color change detection task: ∼13 minutes). Regardless of task, the resulting moving average signal captured changes in the measurement greater than several minutes, which is why we refer to this timescale as minute to minute fluctuations. By subtracting the average of each 100-trial bin from each trial in the bin, the mean-subtracted trial measurements captured changes faster than the bin length, which we refer to as second to second timescale fluctuations. We chose this bin size because it captured fluctuations over tens of minutes that we have explored in prior work linked to impulsivity and engagement (39,60,61). Nevertheless, when we tested other bin sizes for the moving average (20 to 400 trials) for different sessions, we found similar results. Day to day variability in RT was simply estimated by looking at differences from session to session, which were recorded on different days.

### RT variability from second to second

First, we asked how much do RT and HR vary from second to second. Using the values remaining on each trial after subtracting the 100-trial moving average (i.e. the residual values), we estimated second to second variability by computing the standard deviation across these values for each session (σ_s2s,_ Fig 3A,B). In subjects performing a task where difficulty varied from trial to trial (task 2: color change detection task), some of the second to second RT variability could be explained by trial difficulty, where there were slower RTs for trials with more difficult changes (i.e., note the higher overall variability shown in Fig 3A for task 2).

**Fig 3.**
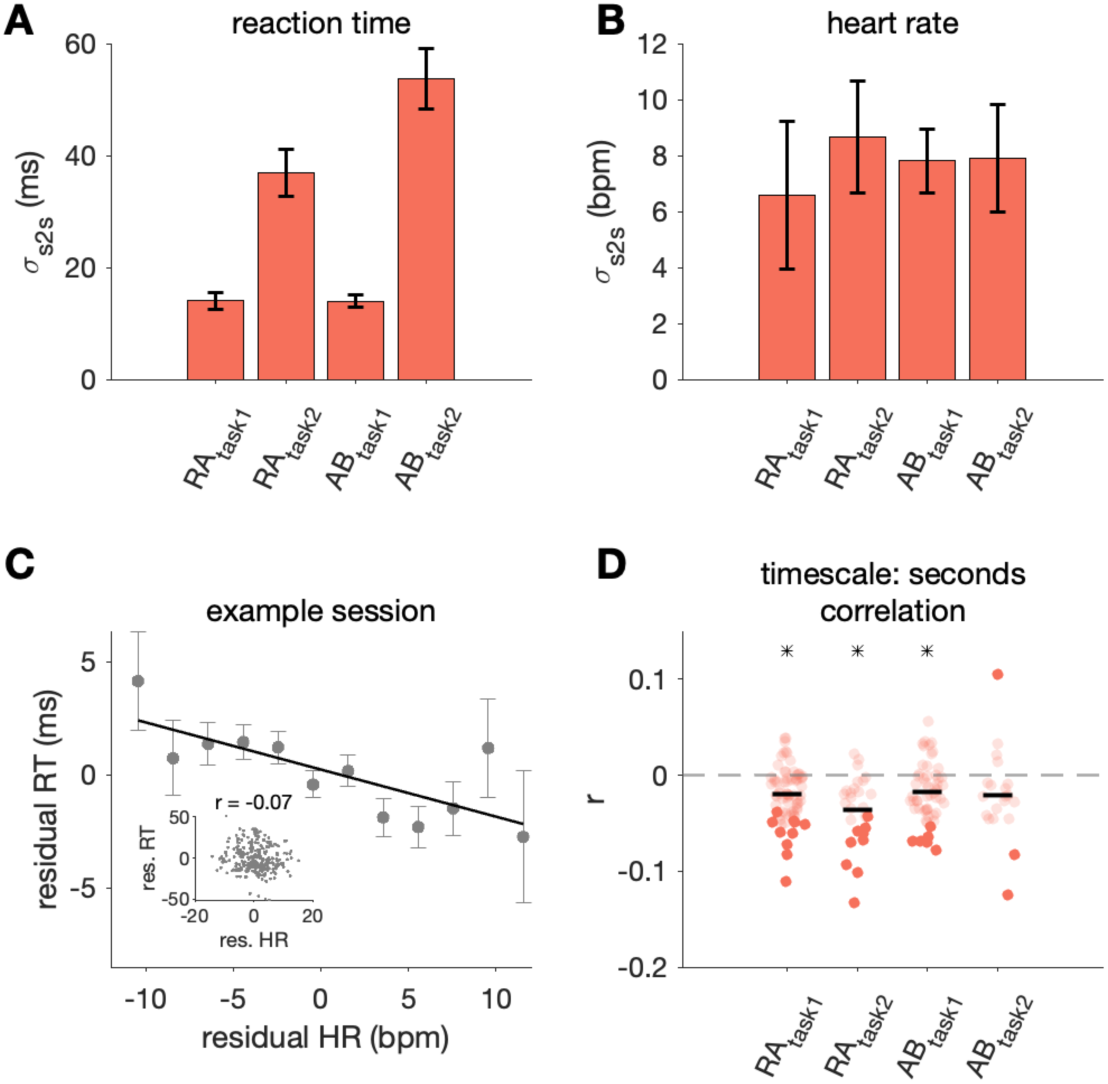
RT and HR varied over seconds. A-B) For each session, *σ_s2s_* was computed as the standard deviation of RT and HR values for each trial (after subtracting the 100-trial moving average from each trial), which captured the second to second fluctuations in these variables (bars show mean and standard deviation of this metric across sessions). C) Second to second (i.e. residual) HR vs. RT for an example session from Subject RA simple saccade task (task 1). Points show mean *±* standard error of RT values for trials within HR bins of 2 bpm for easier visualization of the negative trend. Inset shows second to second HR and RT values for all trials. D) Correlation between second to second HR and RT for all sessions (black line: mean across sessions, asterisk indicates significance across sessions, p< 0.05). Darker points correspond to individual sessions that were significantly different from zero (p < 0.05). Grey outlined point corresponds to the example session in C).

Regardless of the exact source of variability over seconds, we sought to understand how much of the RT variability could be linked to HR. We found second to second RT and HR were negatively correlated in all subjects (Fig 3C and D). For three out of four data sets, the HR-RT correlation across sessions was significantly non-zero, tested using a two tailed t-test with an alpha of 0.05 after Fisher’s r to z transform (RA_task1_: r = −0.02, *p* < 0.0001; RA_task2_: r = −0.04, *p* < 0.0001; AB_task1_: r = −0.02, *p* < 0.0001). In most sessions, when HR was higher, RT was faster. Thus, a significant, albeit small, amount of RT variability could be linked to systemic physiology from second to second. We also performed an analysis to control for task difficulty by using only trials with easy difficulties (difficulties where performance accuracy was typically greater than 90%). As expected, second to second RT variability decreased (RA_task2_ mean σ_s2s_ = 28.8 ms; AB_task2_ mean σ_s2s_ = 33.8 ms). Variability in second to second HR and the correlation between HR and RT remained similar to our observations when using trials of all difficulties.

### RT variability from minute to minute over hundreds of trials

Next, we wanted to understand to what extent RT and HR varied from minute to minute. Figure 4A and C demonstrate how RT and HR drifted from minute to minute (measured by computing average RT and HR in 100-trial bins) for an example session. This drift can be compared to slow fluctuations due to chance by first shuffling the trial order prior to computing binned averages (grey line in Fig 4A and C). Using the 100-trial binned averages, we computed the standard deviation of RT and HR across the binned values for each session (σ_m2m,_ Fig 4B and D). In all cases, except for RT in Subject AB task 2, the variability in RT and HR over minutes was significantly different from that for the shuffled control described above (*p* < 0.05).

**Fig 4.**
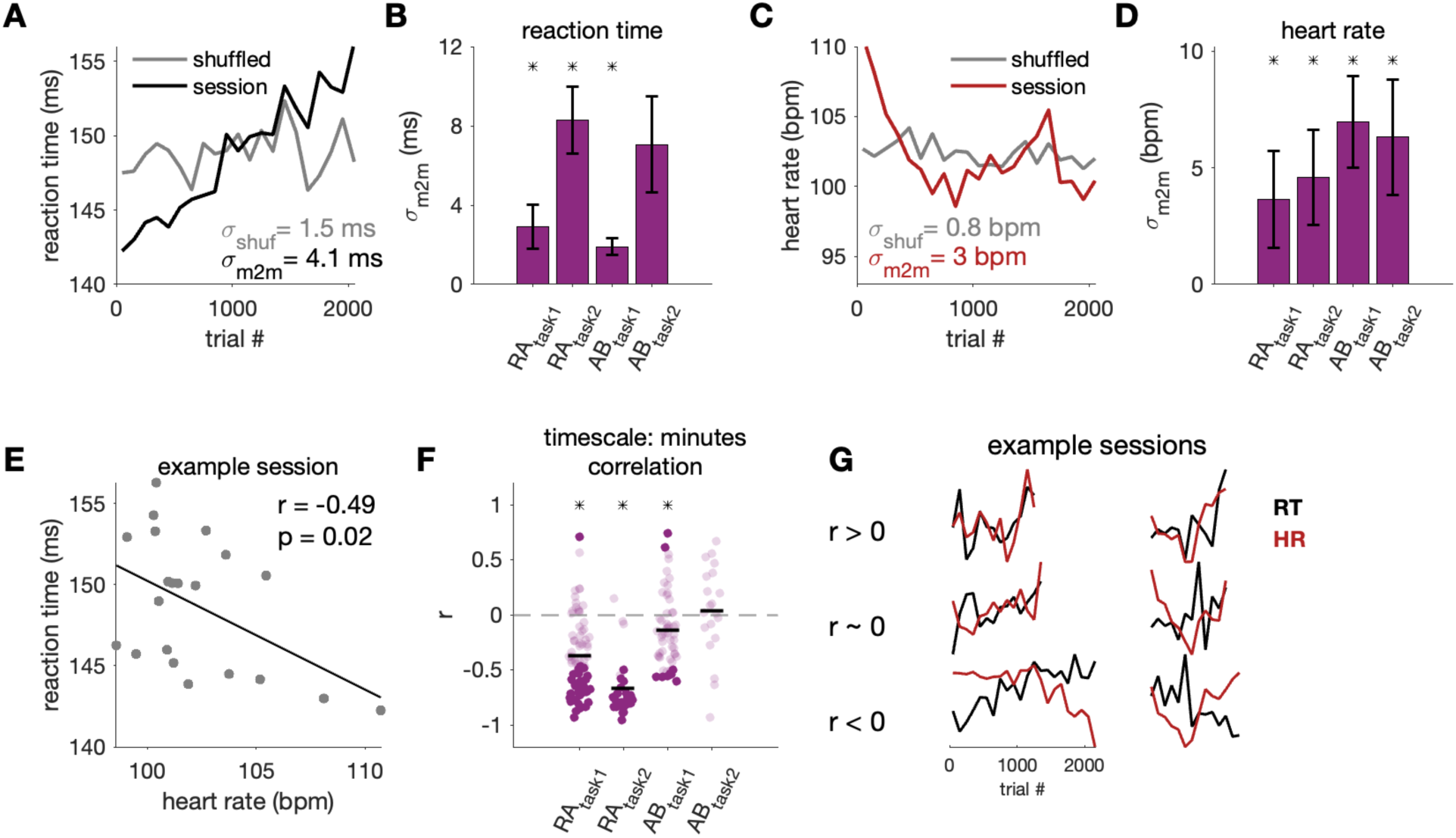
RT and HR varied over minutes. A) 100-trial moving average RT (black) and shuffled control (grey) for one example session, which captured changes slower than tens of minutes. B) Standard deviation of binned RT (*σ_m2m_*). Bars show mean and standard deviation of this metric across sessions. Asterisk indicates subjects where *σ_m2m_* was significantly different from that of the shuffled control (p<0.05). C-D) Same as A-B) but for HR. E) Binned HR vs. binned RT for an example session. F) Correlation between binned HR and RT (black line: mean across sessions; asterisk indicates significance across sessions, p< 0.05). Darker points correspond to individual sessions that were significantly different from zero (p < 0.05). G) Additional example sessions demonstrating the variability in the relationship between HR and RT across sessions.

The minute to minute RT drift was also significantly negatively correlated with minute to minute fluctuations in HR (Fig 4E and F). For three out of four data sets, the minute to minute HR-RT correlation across sessions was significantly non-zero (Ra_task1_: r = −0.37, *p* < 0.0001; Ra_task2_: r = −0.67, *p* < 0.0001; Ab_task1_: r = −0.14, *p* = 0.004; Ab_task2_: r = 0.04, *p* = 0.96). Generally, when HR trended higher over hundreds of trials, RT was also faster. However, there were individual sessions in all subjects where HR and RT had the opposite relationship (Fig 4G). These findings were robust to different choices in the number of trials per bin (ranging from 20 to 400 trials) to produce the minute to minute average traces.

Given the variability in the sign of the relationship between HR and RT from session to session, we assessed whether the direction of the relationship was consistent over seconds and over minutes for the same session. In other words, for a given session, when there was a positive relationship between HR and RT over seconds was there also a positive relationship between HR and RT over minutes. We found the sign of the HR-RT relationship was consistent over seconds and over minutes for the majority of sessions (65.3% of RA_task1_ sessions, 77.8% of RA_task2_ sessions, 64.8% of AB_task1_ sessions, and 55.0% of AB_task2_ sessions).

### HR explained variance in RT beyond session time effects

There are many reasons why RT and HR may trend together over minutes across the session. One explanation is simply the passage of time during the session as the subject gradually becomes fatigued or satiated. We asked whether HR explained additional variance in RT beyond session time effects. We performed a multiple linear regression to predict RT using the bin number (a proxy for session time) and HR (R^2^ for RA_task1_: 0.39, RA_task2_: 0.64, AB_task1_: 0.03, AB_task2_: 0.11). To create a model that could use data from different sessions, we z-scored RT and HR within a session. To assess whether HR added additional information to predicting RT, we created a null distribution by computing R^2^ for a linear regression fit using the bin numbers and shuffled HR values and repeated this 1000 times. The p-value was calculated as the proportion of bootstrapped null R^2^ values that were greater than or equal to the true R^2^ value since the null hypothesis was that HR did not explain additional variance. For subject RA in task 1 and 2, and for subject AB in task 1, the true R^2^ was significantly different from the shuffled null distribution, which meant HR explained additional variance beyond time in session (fraction of null R^2^ >= data R^2^ - RA_task1_: 0/1000, RA_task2_: 0/1000, AB_task1_: 0/1000, AB_task2_: 638/1000).

We considered whether the significant relationship between HR and RT (Fig 3D, 4F) might be driven by the longest or shortest sessions. To control for this, we removed 20% of sessions that were the longest in session length as well as 20% of sessions that were the shortest in session length. We found that the HR and RT correlations were similar to the full data when we removed either the shortest or longest sessions from consideration.

### RT variability between sessions across days

In addition to variability within a session from second to second and minute to minute, we wanted to quantify the extent to which RT and HR varied on longer timescales. We therefore assessed how much the subject’s RT and HR changed from day to day despite performing the same, well-practiced task. To estimate this variability we created a distribution of session-averaged RTs. Since each session was a different length, we used the mean of a subset of trials from the beginning of each session to capture a comparable part of the subject’s task behavior from day to day (mean of correct trials 200-300; note we excluded the first 200 trials to allow behavior to stabilize and reduce the impact of variability we observed in the start of the session on our analysis). This gave us a single representative RT value for each session to create the across session distribution that consisted of N values (N = # of sessions). We also created a null distribution that included all trials from all sessions. We sought to compute a standard deviation metric similar to σ_m2m_ used in Figure 4. First, we found the number of bins in each session (ex. L_1_ for session 1, L_2_ for session 2, etc.). Then, we grabbed L_1_ values from the across session distribution and computed the standard deviation across these values (σ_d2d,1_). We also grabbed L_1_ values from the null distribution (σ_shuff,1_). We repeated this for all values of L (i.e. the number of bins per session for each session). This gave us N σ_d2d_ and σ_shuff_ values which allowed us to perform a t-test to compare the day to day variability in the data and the shuffled control. We repeated this procedure for HR (Fig 5A-D) and found that the day to day variability in the RT and HR data were significantly different from that of the shuffled controls for all subjects (*p* < 0.05). This indicates there was substantial day to day variability in RT in both of our subjects performing both tasks.

**Fig 5.**
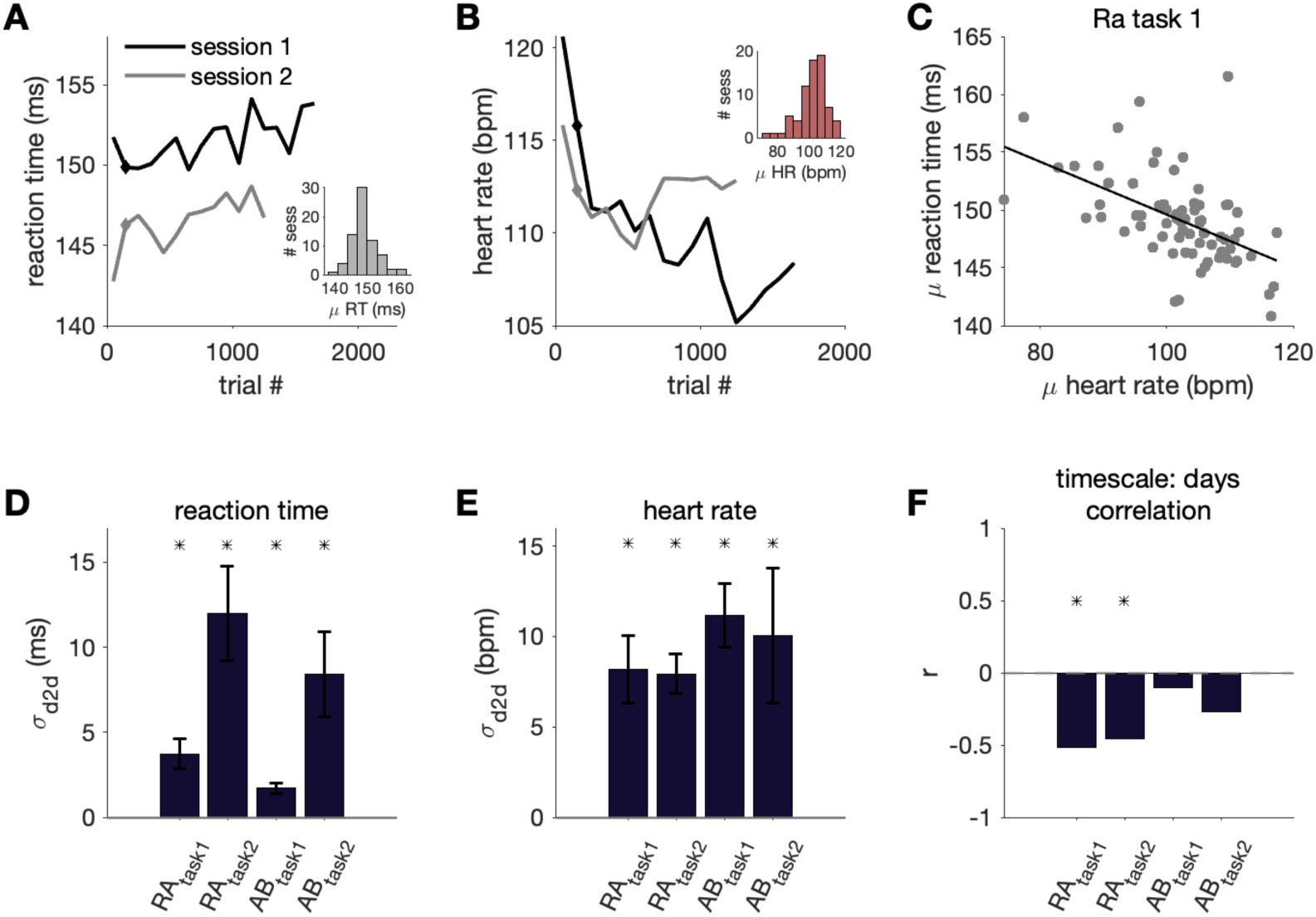
RT and HR varied across sessions. A) RT over the session for two example sessions (100-trial binned moving average). We used the mean of trials 200 to 300 for each session as a representative RT value (indicated with a diamond point). The inset shows the distribution of these mean RT values across all sessions for subject RA in task 1. D) Standard deviation of the RTs across sessions (*σ_d2d_*) for each subject (bars show mean and standard deviation of this metric). B,E) Same as A,D) but for HR. C) Mean session HR vs. mean session RT for all sessions for subject RA in the simple saccade task (black line: line of best fit). F) Correlation between mean session HR and RT for each subject. Asterisk indicates the correlation value was significantly different from the shuffled control (p<0.05).

Furthermore, in subject RA across two different tasks, the session HR was significantly correlated with the session RT (Fig 5E and D; RA_task1_: r = −0.52, *p* < 0.0001; RA_task2_: r = −0.45, *p* = 0.02). This indicates there was a link between day to day variability in RT and HR: on days the average HR was higher, the average RT was also faster. Although there was a similar direction of HR-RT trends in subject AB across two different tasks, the relationship was not statistically significant (AB_task1_: r = −0.10, *p* = 0.46; AB_task2_: r = −0.27, *p* = 0.25).

### Variability in RT and HR controlled for time of day

Some of the day to day variability we observed could be due to circadian factors that lead to differences in performance across time of day. Although it was typical for our subjects to be run at a relatively consistent time of day, we did not explicitly control this in the main data sets. Therefore, in a subset of sessions we attempted to control for factors related to time of day. AM sessions were recorded at the same time of day each day on consecutive days (number of AM sessions: RA_task2_ - 6; AB_task1_ - 6). PM sessions were recorded on a different set of consecutive days in a similar manner (number of PM sessions: RA_task2_ - 12; AB_task1_ - 10). This helped to control for circadian-related fluctuations in behavior and physiology as well as other factors linked to satiety and fatigue based on the time since last task performance (on the previous day) and scheduled feedings.

For each session, we computed a single representative RT and HR value using the mean of trials 200 to 300 as in Figure 5. In the table below, σ_d2d_ was computed as the standard deviation across both AM and PM sessions. σ_AM_ and σ_PM_ were computed as the standard deviation across AM only or PM only sessions respectively (Table 1). If time of day explained a large amount of variability from session to session, then we would expect σ_AM_ and σ_PM_ to be lower than the standard deviation across all sessions combined (σ_d2d_). We found the standard deviation when controlling for time of day was lower than the standard deviation across all sessions combined (binomial exact test: *p* = 0.04), indicating that some of the day to day variation we observed was due to circadian fluctuations. Nevertheless, while time of day explained some of the variability, there was still a large amount of day to day variability even for sessions recorded at the exact same time of day.

**Table 1.** Variability in RT and HR for AM and PM sessions. Standard deviation of the RTs and HRs across combined AM and PM sessions (*σ_d2d_*) for a subset of sessions in Ab and Ra. *σ_AM_* was computed as the standard deviation of RTs and HRs across AM sessions only. *σ_PM_* was computed as the standard deviation of RTs and HRs across PM sessions only. The last column shows the absolute value of the difference between the mean of the AM and PM sessions. Instances where the variability was reduced by separating AM and PM sessions are in bold.

| | | $\sigma_{d2d}$ | $\sigma_{AM}$ | $\sigma_{PM}$ | $\vert \mu_{AM} - \mu_{PM}\vert $ |
| --- | --- | --- | --- | --- | --- |
| RT | Ab<sub>task1</sub> | 1.8 ms | **1.6 ms** | **1.3 ms** | 1.6 ms |
|  | Ra<sub>task2</sub> | 12.4 ms | **6.2 ms** | **8.7 ms** | 12.1 ms |
| HR | Ab<sub>task1</sub> | 10.5 bpm | **6.3 bpm** | **8.4 bpm** | 4.8 bpm |
|  | Ra<sub>task2</sub> | 8.2 bpm | 9.0 bpm | **7.4 bpm** | 4.9 bpm |

### The relationship between linked measures of arousal and behavior is complex

Thus far, we have used HR as a proxy for arousal. We wanted to assess how well HR aligned with perhaps the most commonly used physiologic proxy for arousal: pupil diameter (Fig 6A). For each trial, we computed mean pupil diameter during the delay period (200-400 ms post fixation). We computed correlation between pupil diameter and HR on both the second to second timescale and on the minute to minute timescale. The minute to minute timescale for pupil and HR was the 100-trial moving average, as in Figure 4, while the second to second timescale was computed by subtracting the 100-trial bin average from each trial in that bin, as in Figure 3.

**Fig 6.**
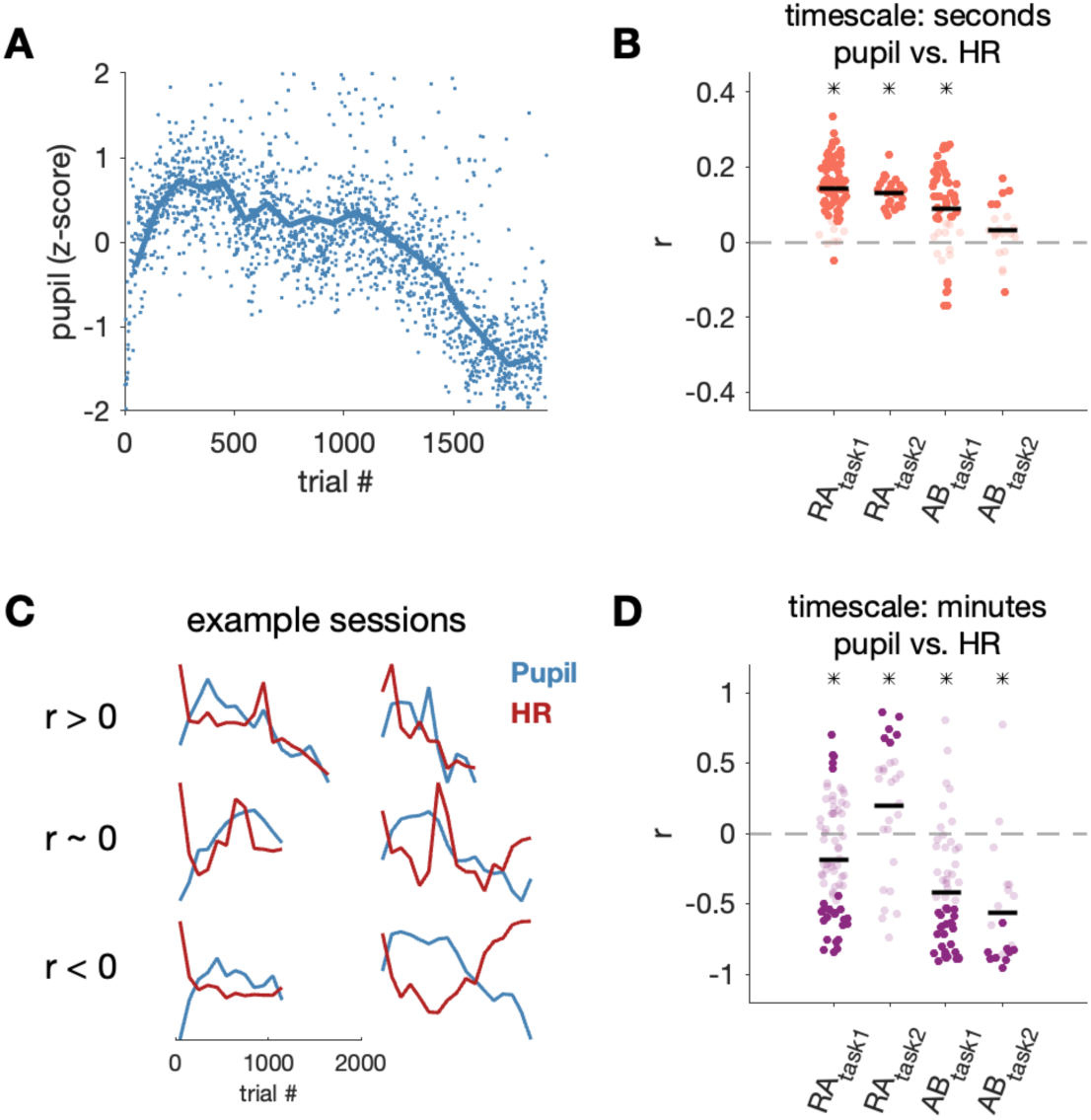
HR was linked to pupil in a timescale-dependent way. A) Z-scored pupil diameter values for all correct trials over time, excluding points with pupil values greater than 2 standard deviations from the mean (blue line: moving average using 100-trial bins). B) Correlation between residual pupil and HR on every trial. C) Additional example sessions showing the relationship between 100-trial binned averages of pupil diameter and HR. D) Correlation between binned pupil and HR (black line: mean across sessions; asterisk indicates significance across sessions, p< 0.05). Darker points correspond to individual sessions that were significantly different from zero (p < 0.05).

From second to second, HR and pupil diameter were significantly correlated in three out of four subjects (Fig 6B; RA_task1_: r = 0.14, *p* < 0.0001; RA_task2_: r = 0.13, *p* < 0.0001; AB_task1_: r = 0.09, *p* < 0.0001). This indicated that when HR was higher on a trial, the pupil diameter was usually also larger on that trial. Interestingly, the opposite was true on a slower timescale of minutes (Fig 6D). In three out of four data sets, we found slow changes in pupil and HR were significantly negatively correlated (RA_task1_: r = −0.19, *p* = 0.0001; AB_task1_: r = −0.42, *p* < 0.0001; AB_task2_: r = −0.56, *p* < 0.0001). Thus, when HR was higher over several minutes, pupil diameter tended to be smaller on the same timescale. One subject on task 2 (color change detection task) had a significant positive correlation between pupil diameter and HR from minute to minute (RA_task2_: r = 0.20, *p* = 0.04), and there were individual sessions in each subject that showed a positive relationship (Fig 6C and D).

Given the timescale-dependent relationship between HR and pupil, it is possible that the relationship between arousal-linked physiology and behavior may depend on the physiology measure to which behavior is compared. We also characterized the relationship between pupil diameter and RT for both the fast timescale over seconds (Fig 7A) and the slow timescale over minutes (Fig 7B). If the direction of the RT-pupil relationship was the same as the direction of RT-HR relationship, it supports the commonly held assumption that different biomarkers of arousal have a consistent relationship with behavior. In other words, fast HR and a large pupil diatmeter, which are both linked to mechanisms of increased arousal, have a consistent relationship with RT. In contrast, if the directions of the RT-pupil and RT-HR relationships were different, it suggests the link between different biomarkers of arousal and behavior may be more complex.

**Fig 7.**
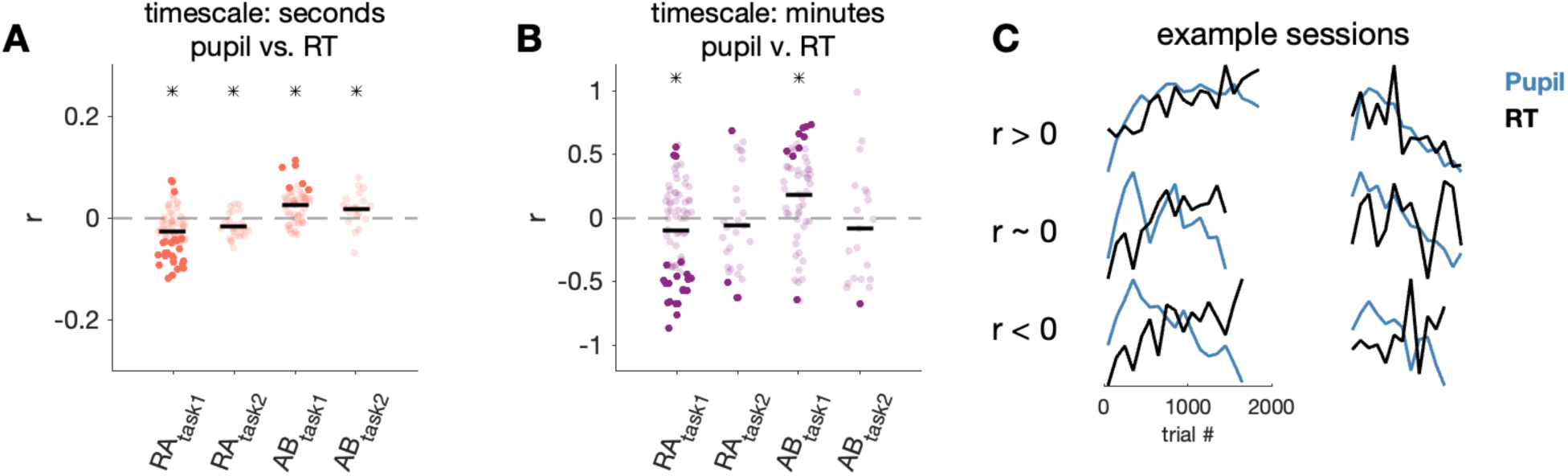
RT was weakly linked to pupil. A) Correlation between residual pupil and RT on every trial (black line: mean across sessions; asterisk indicates significance across sessions, p< 0.05). Darker points correspond to individual sessions that were significantly different from zero (p < 0.05).. B) Correlation between 100-trial binned averages of pupil and HR. C) Additional example sessions showing the relationship between binned averages of pupil diameter and RT.

The RT-pupil relationship varied substantially across sessions, as can be seen by the large spread of correlation values across sessions as well as by observing individual example sessions (Fig 7C). In one subject, across both task types and timescales there was an average negative correlation between pupil and RT (subject RA). This indicates faster RTs were associated with larger pupil diameters, and mirrors the general negative HR-RT relationship. However, in the second subject, there was an average positive correlation between pupil and RT on the second to second timescale and, in one task, on the minute to minute timescale (subject AB). This trend was in the opposite direction of the general HR-RT relationship, which points towards a more complex relationship between different measures of physiology and behavior.

We also analyzed additional RT and pupil data from two subjects (subject PE and subject WA) in previously published data (*60*). In that dataset, subjects performed a memory guided saccade task, where they had to remember and look to the location of a briefly presented target, and an orientation change detection task, where they had to detect a change in orientation of grating stimuli. Similar to subject RA, subjects PE and WA generally showed an average negative RT-pupil correlation on both timescales across sessions but also had many sessions with a positive RT-pupil correlation (Table 2). Thus, in aggregate across sessions and subjects there was generally a consistent direction of RT-HR and RT-pupil relationships and therefore, overall (but not absolute) consistency in the different measures of arousal and behavior.

**Table 2.** Relationship between RT and pupil for two additional subjects from Johnston et al (*60*). Asterisk indicates significance across sessions (p < 0.05).

|  |  | correlation (r) |  |
| --- | --- | --- | --- |
|  |  | timescale: seconds | timescale: minutes |
| Subject PE | Memory guided saccade (N = 31) | -0.01 | -0.29* |
|  | Orientation change detection (N = 24) | -0.07* | -0.21 |
| Subject WA | Memory guided saccade (N = 21) | -0.07* | -0.47* |
|  | Orientation change detection (N = 23) | 0.01 | 0.03 |

Given that on average RT was negatively related to both pupil diameter (Fig 7B) and HR (Fig 4F) over a timescale of minutes, one might expect pupil diameter and RT to have a positive relationship with each other. However, pupil diameter and HR were negatively correlated from minute to minute (Fig 6D). This could indicate there is one component of HR linked to pupil diameter and a separate component of HR linked to behavior. To quantitatively assess this, we asked whether the relationship between RT and HR changed when controlling for pupil diameter using partial correlation analysis. If the relationship between RT and HR persisted (i.e. was non-zero), then it cannot be completely explained by a common link with pupil. We found that even when accounting for variance explained by pupil diameter, the partial correlation between RT and HR was negative in most subjects on timescales of seconds and minutes (Table 3). Thus, the correlation between HR, pupil, and RT was not simply explained by a simple linear relationship between all three measurements.

**Table 3.** Partial correlation between RT and HR across session controlling for pupil diameter. Asterisk indicates significance across sessions (p < 0.05).

| | | partial correlation ($\rho$) | |
| --- | --- | --- | --- |
|  |  | timescale:<br>seconds | timescale:<br>minutes |
| Subject RA | Task 1 | -0.01* | -0.41* |
|  | Task 2 | -0.03* | -0.64* |
| Subject AB | Task 1 | -0.02* | -0.04 |
|  | Task 2 | 0.02 | 0.03 |

In summary, there was a large degree of across session and across subject variability in the relationship between different measures of arousal and behavior. First, we found that HR and pupil diameter, which are both independently tied to physiological arousal mechanisms, had a predictable positive relationship over seconds (fast HR, large pupil), but a counterintuitive negative relationship over many minutes (fast HR, small pupil). Despite this surprising HR-pupil relationship, in general both HR and pupil diameter had a negative relationship with RT. Additionally, the relationship between HR and RT could not be accounted for by a linear relationship of each variable with pupil diameter. Together, these results suggest RT is linked to both HR and pupil diameter, albeit with different strengths and not due to a single common relationship between all three measurements. This could be consistent with HR and pupil partially indexing separate but connected arousal systems.

## DISCUSSION

Even in a laboratory environment with controlled conditions, daily experiments, and highly trained subjects, there is a large amount of variability in RT. Some of this variability can be attributed to underlying arousal processes (18,19,26,33). In studies that lack a physiological arousal measure, such as HR or pupil diameter, RT has been used as an indicator of arousal state, where faster RTs are inferred to correspond to a higher arousal level in which the subject is more ready to react. To strengthen this interpretation, studies that characterize the link between RT and physiological arousal measures are critical. We quantified the variation in RT and HR across different timescales (Fig 8) and found there was a significant relationship between them from second to second, minute to minute, and day to day.

**Fig 8.**
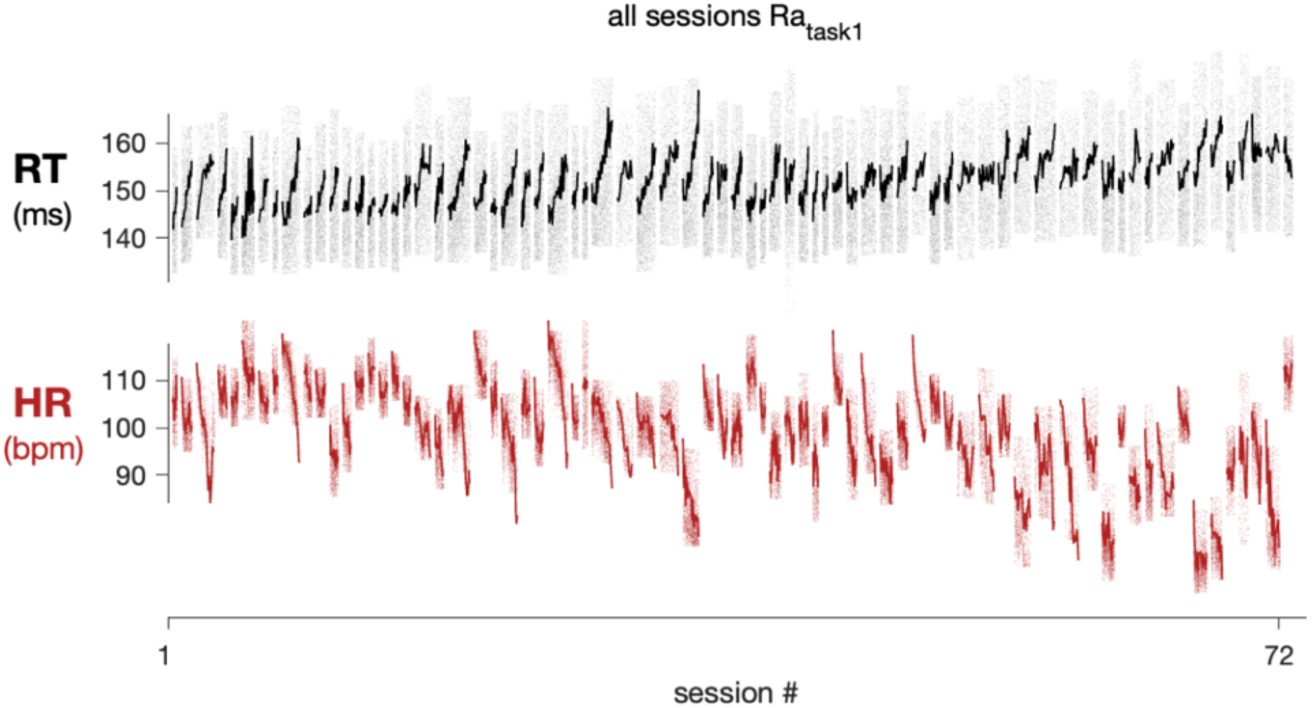
RT and HR varied on timescales of seconds (black/red points), minutes (black/red lines), and days (every session). RT and HR for all sessions (N=72) for Subject RA task 1 (simple saccade task). Sessions are ordered from least to greatest mean RT (computed using trials 200-300 on each session).

### Relating RT to a physiological marker of arousal

In general, we expected an elevated arousal level to correspond to a higher HR due to activation of the sympathetic nervous system (62). Given that faster RTs are also linked to elevated arousal, we predicted a negative correlation between HR and RT (higher HRs and lower RTs), which we observed across subjects and tasks on timescales over seconds and minutes. On a timescale over days, in one subject, there was also a significant negative correlation, while in the second subject the relationship also trended negative but was not significant. Interestingly, there were numerous sessions where HR and RT had a positive relationship, particularly on the minute to minute timescale. Although there are a few examples in the literature with a positive HR-RT correlation (53), our general intuition regarding the negative direction of the relationship between HR and RT comes from large, evoked changes, such as in response to an alarming stimulus (63,64). For example, if we suddenly come across a snake on a hiking trail, our HR would increase and RT might decrease as we enter a ‘fight or flight’ state. As a result, we would observe a strong negative linear relationship between HR and RT from the pre- to post-snake time periods. We have less intuition about these relationships in everyday circumstances, such as during continued performance of a well-known task. By characterizing the HR-RT relationship under these conditions, we can better understand how to interpret more typical arousal fluctuations that are weaker than those evoked by a sudden, frightening stimulus like a snake.

### Sources of variability

The large amount of variability in the strength and magnitude of the HR-RT correlation is likely indicative of other influences on HR and RT that become more dominant on certain timescales and in different sessions. For example, from second to second, changes in physiology such as blood pressure (65) or respiration (66) may have a larger influence on HR than processes that are linked to RT (Fig 9). Similarly, from second to second, subtle changes in muscle tone might affect RT (67) but have no relationship to arousal changes.

**Fig 9.**
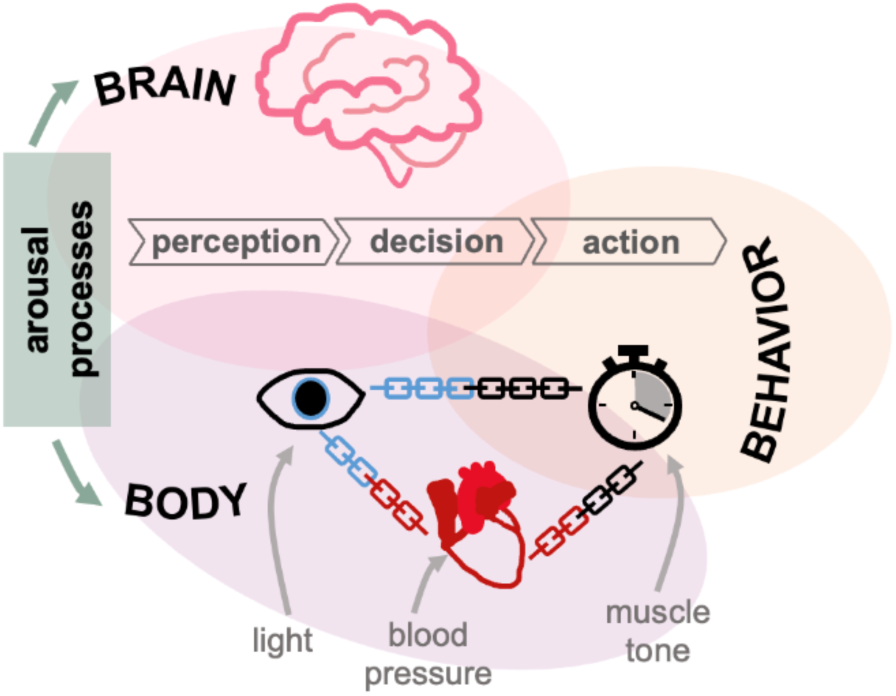
Sources of RT and physiology variability. Schematic demonstrating how multiple arousal processes might affect both the brain and body physiology, such as HR and pupil. Dark grey boxes (light, BP-blood pressure, muscle) indicate processes that may independently influence certain measurements. Light grey boxes (perception, decision, action) reflect brain processes that determine the behavioral outcome.

The strong correlations we observed over minutes between HR and RT could have arisen from slow arousal-linked processes that drift over time, such as fatigue, satiation, and engagement. Time since the session began (session time) can be used to approximate a combination of fatigue and satiation since the subjects grow more tired and earn more liquid reward as a function of time. As expected, session time explained a significant amount of the variability in RT. However, even after accounting for this time-related variability, HR explained a significant amount of RT variability. This suggests the relationship between RT and HR cannot be solely explained by a simple process that ramps up or down over the course of the session. In an ideal experiment, we could manipulate arousal level directly and then measure the effect on RT and HR. A significant challenge when studying arousal is that even when controlling for every physical variable (level of thirst, quantity of sleep, etc.) it is difficult, if not impossible, to control emotional state and temperament, which also influence arousal level and behavior (51,68–70). For example, imagine a scenario immediately prior to the start of the session where our monkey subject was engaged in a charged social interaction with another member of their colony. They may be more impulsive due to a heightened emotional state, resulting in overall faster RTs. Since an arousal-related emotional process is driving their behavior, there may be a strong relationship between HR and RT. In comparison, a monkey who begins a session in a more neutral emotional state due to the lack of a prior antagonistic interaction may be more susceptible to independent influences on their HR and RT than common arousal processes. Similarly, a subject with a calmer baseline temperament may be less driven by common arousal processes than one with a more volatile temper. These factors could be a few possible sources of the inter-subject variability we observed in the HR-RT relationship between Subject AB and RA.

### Are all biomarkers of arousal equivalent?

With some exceptions (49,50,71), it is rare for studies to measure multiple physiological markers of arousal simultaneously. Particularly in cognitive studies, pupil diameter is the most common measure of global arousal processes due to its partial control by a core brainstem arousal region (46,47,72–74) and due to prior work that demonstrated a link between pupil size and RT (56,57). It may be assumed that physiological metrics will be highly correlated with each other and by extension arise from a common underlying arousal process. Driven by the sympathetic nervous system, we would predict that an increased HR would occur with an increased pupil diameter. While we observed this positive relationship on fast timescales over seconds, on average we observed the opposite on slower timescales over minutes in three out of four data sets. In these data sets, there were still many sessions that had the predicted positive relationship between HR and pupil diameter; however, the majority had a negative relationship. Beyond measurement noise that could also vary from session to session, this inverted relationship could be driven by independent sources of variability that drive one physiological metric but not the other (Fig 9) or different baseline emotional states when starting the session (as described above).

Another possibility could be there are multiple underlying arousal processes that operate on different timescales (over seconds and over minutes) and affect HR and pupil diameter differently. Locus coeruleus activity has been linked to changes in both pupil diameter (47) and HR (75). However, HR is also influenced by another brainstem nucleus called the nucleus ambiguus (42). These brain regions have direct and indirect connections to each other as well as to numerous other brainstem regions controlling the autonomic nervous system (42,76). It is possible that there are some arousal processes in which these brain regions act in coordination to influence HR, pupil diameter, and RT, while there are also other processes where they act on only one or two of these variables. Multiple arousal processes acting simultaneously could explain how the negative relationships between RT and both pupil and HR could co-occur with the negative relationship between HR and pupil diameter over minutes. Although we can only speculate the source of the negative HR-pupil diameter relationship, it is clear that HR and pupil diameter should not be considered interchangeable markers of the same global arousal processes under all circumstances.

One method of further teasing apart these processes in future work would be to separate the different components of RT using a model such as drift diffusion or a linear ballistic accumulator (1,2,10). One could then ask whether the components that influence RT, such as the drift rate, starting point, and boundary separation, are the ones that are linked to HR and pupil. Additionally, we chose to quantify the relationship between RT and arousal biomarkers using Pearson’s correlation coefficient because it is commonly used throughout the literature and easily interpretable. However, correlation does not characterize the non-linear relationships that may be present between these variables. Future work may consider how behavior, physiology, and arousal processes can be modeled non-linearly (77). This would permit models in which the direction of the relationship between variables, such as HR and RT, varies over the session, one possible phenomenon that could give rise to weak correlations when constrained to a linear relationship framework. Nevertheless, our work supports the conclusion that multiple arousal mechanisms can act simultaneously to influence behavior across timescales. It also emphasizes the utility of simultaneously measuring multiple physiological signals to better isolate arousal processes that give rise to different aspects of behavior.

## ACKNOWLEDGEMENTS

D.I. was supported by NIH F30 MH129056, the Ronald F. and Janice A. Zollo Fellowship, the Carnegie Mellon University Neuroscience Institute Carnegie Fellowship, and the ARCS Foundation Thomas-Pittsburgh Chapter award. M.A.S., J.M.K., and M.R.G were supported by NIH R01 MH128393. E.E.S. was supported by NIH T32 EB029365 and the Ronald F. and Janice A. Zollo Fellowship. We would also like to thank Samantha Nelson for her technical and experimental support as well as our animal care staff.

## REFERENCES

1. Ratcliff R. Parameter variability and distributional assumptions in the diffusion model. Psychol Rev. 2013;120(1):281–92. doi:10.1037/a0030775

2. Ratcliff R, McKoon G. The diffusion decision model: theory and data for two-choice decision tasks. Neural Comput. 2008 Apr;20(4):873–922. doi:10.1162/neco.2008.12-06-420 PubMed PMID: 18085991; PubMed Central PMCID: PMC2474742.

3. Smith PL. Psychophysically principled models of visual simple reaction time. Psychol Rev. 1995 Jul;102(3):567–93. doi:10.1037/0033-295X.102.3.567

4. Schmolesky MT, Wang Y, Hanes DP, Thompson KG, Leutgeb S, Schall JD, et al. Signal timing across the macaque visual system. J Neurophysiol. 1998 Jun;79(6):3272–8. doi:10.1152/jn.1998.79.6.3272 PubMed PMID: 9636126.

5. Maunsell JH, Gibson JR. Visual response latencies in striate cortex of the macaque monkey. J Neurophysiol. 1992 Oct;68(4):1332–44. doi:10.1152/jn.1992.68.4.1332 PubMed PMID: 1432087.

6. Sparks DL. The brainstem control of saccadic eye movements. Nat Rev Neurosci. 2002 Dec;3(12):952–64. doi:10.1038/nrn986 PubMed PMID: 12461552.

7. Bruce CJ, Goldberg ME, Bushnell MC, Stanton GB. Primate frontal eye fields. II. Physiological and anatomical correlates of electrically evoked eye movements. J Neurophysiol. 1985 Sep 1;54(3):714–34. doi:10.1152/jn.1985.54.3.714

8. Cheney PD, Fetz EE. Functional classes of primate corticomotoneuronal cells and their relation to active force. J Neurophysiol. 1980 Oct 1;44(4):773–91. doi:10.1152/jn.1980.44.4.773

9. Morrow MM, Miller LE. Prediction of Muscle Activity by Populations of Sequentially Recorded Primary Motor Cortex Neurons. J Neurophysiol. 2003 Apr 1;89(4):2279–88. doi:10.1152/jn.00632.2002

10. Brown SD, Heathcote A. The simplest complete model of choice response time: linear ballistic accumulation. Cognit Psychol. 2008 Nov;57(3):153–78. doi:10.1016/j.cogpsych.2007.12.002 PubMed PMID: 18243170.

11. Fengler A, Bera K, Pedersen ML, Frank MJ. Beyond Drift Diffusion Models: Fitting a Broad Class of Decision and Reinforcement Learning Models with HDDM. J Cogn Neurosci. 2022 Sep 1;34(10):1780–805. doi:10.1162/jocn_a_01902

12. Noorani I, Carpenter RHS. The LATER model of reaction time and decision. Neurosci Biobehav Rev. 2016 May;64:229–51. doi:10.1016/j.neubiorev.2016.02.018

13. Gold JI, Shadlen MN. The neural basis of decision making. Annu Rev Neurosci. 2007;30:535–74. doi:10.1146/annurev.neuro.29.051605.113038 PubMed PMID: 17600525.

14. Palmer J, Huk AC, Shadlen MN. The effect of stimulus strength on the speed and accuracy of a perceptual decision. J Vis. 2005 May 2;5(5):1. doi:10.1167/5.5.1

15. Lo CC, Wang XJ. Cortico–basal ganglia circuit mechanism for a decision threshold in reaction time tasks. Nat Neurosci. 2006 Jul;9(7):956–63. doi:10.1038/nn1722

16. Hanes DP, Schall JD. Neural control of voluntary movement initiation. Science. 1996 Oct 18;274(5286):427–30. doi:10.1126/science.274.5286.427 PubMed PMID: 8832893.

17. Jensen AR. The importance of intraindividual variation in reaction time. Personal Individ Differ. 1992 Aug;13(8):869–81. doi:10.1016/0191-8869(92)90004-9

18. Guan S, Liu Y, Xia R, Zhang M. Covert attention regulates saccadic reaction time by routing between different visual-oculomotor pathways. J Neurophysiol. 2012 Mar 15;107(6):1748–55. doi:10.1152/jn.00082.2011

19. Yamashita A, Rothlein D, Kucyi A, Valera EM, Germine L, Wilmer J, et al. Variable rather than extreme slow reaction times distinguish brain states during sustained attention. Sci Rep. 2021 Jul 21;11(1):14883. doi:10.1038/s41598-021-94161-0 PubMed PMID: 34290318; PubMed Central PMCID: PMC8295386.

20. Posner MI, Nissen MJ, Ogden WC. Attended and unattended processing modes: The role of set for spatial location. In: Modes of perceiving and processing information. 1978. p. 137–57.

21. Summerside EM, Shadmehr R, Ahmed AA. Vigor of reaching movements: reward discounts the cost of effort. J Neurophysiol. 2018 Jun 1;119(6):2347–57. doi:10.1152/jn.00872.2017 PubMed PMID: 29537911; PubMed Central PMCID: PMC6734091.

22. Manohar SG, Chong TTJ, Apps MAJ, Batla A, Stamelou M, Jarman PR, et al. Reward Pays the Cost of Noise Reduction in Motor and Cognitive Control. Curr Biol CB. 2015 Jun 29;25(13):1707–16. doi:10.1016/j.cub.2015.05.038 PubMed PMID: 26096975; PubMed Central PMCID: PMC4557747.

23. Lloyd B, Nieuwenhuis S. The effect of reward-induced arousal on the success and precision of episodic memory retrieval. Sci Rep. 2024 Jan 24;14(1):2105. doi:10.1038/s41598-024-52486-6 PubMed PMID: 38267573; PubMed Central PMCID: PMC10808342.

24. Smoulder AL, Pavlovsky NP, Marino PJ, Degenhart AD, McClain NT, Batista AP, et al. Monkeys exhibit a paradoxical decrease in performance in high-stakes scenarios. Proc Natl Acad Sci. 2021 Aug 31;118(35):e2109643118. doi:10.1073/pnas.2109643118

25. Algermissen J, Den Ouden HEM. High stakes slow responding, but do not help overcome Pavlovian biases in humans. Learn Mem. 2024 Aug;31(8):a054017. doi:10.1101/lm.054017.124

26. Reddi BA, Carpenter RH. The influence of urgency on decision time. Nat Neurosci. 2000 Aug;3(8):827–30. doi:10.1038/77739 PubMed PMID: 10903577.

27. Cisek P, Puskas GA, El-Murr S. Decisions in Changing Conditions: The Urgency-Gating Model. J Neurosci. 2009 Sep 16;29(37):11560–71. doi:10.1523/JNEUROSCI.1844-09.2009

28. Ferrucci L, Genovesio A, Marcos E. The importance of urgency in decision making based on dynamic information. Van Vugt MK, editor. PLOS Comput Biol. 2021 Oct 4;17(10):e1009455. doi:10.1371/journal.pcbi.1009455

29. Mackworth NH. The Breakdown of Vigilance during Prolonged Visual Search. Q J Exp Psychol. 1948 Apr;1(1):6–21. doi:10.1080/17470214808416738

30. Helton WS, Warm JS. Signal salience and the mindlessness theory of vigilance. Acta Psychol (Amst). 2008 Sep;129(1):18–25. doi:10.1016/j.actpsy.2008.04.002

31. Oken BS, Salinsky MC, Elsas SM. Vigilance, alertness, or sustained attention: physiological basis and measurement. Clin Neurophysiol. 2006 Sep;117(9):1885–901. doi:10.1016/j.clinph.2006.01.017

32. Boksem MAS, Meijman TF, Lorist MM. Effects of mental fatigue on attention: an ERP study. Brain Res Cogn Brain Res. 2005 Sep;25(1):107–16. doi:10.1016/j.cogbrainres.2005.04.011 PubMed PMID: 15913965.

33. Soto-Leon V, Alonso-Bonilla C, Peinado-Palomino D, Torres-Pareja M, Mendoza-Laiz N, Mordillo-Mateos L, et al. Effects of fatigue induced by repetitive movements and isometric tasks on reaction time. Hum Mov Sci. 2020 Oct;73:102679. doi:10.1016/j.humov.2020.102679 PubMed PMID: 32980590.

34. Wright KP, Hull JT, Czeisler CA. Relationship between alertness, performance, and body temperature in humans. Am J Physiol-Regul Integr Comp Physiol. 2002 Dec 1;283(6):R1370–7. doi:10.1152/ajpregu.00205.2002

35. Chandrakumar D, Dorrian J, Banks S, Keage H a. D, Coussens S, Gupta C, et al. The relationship between alertness and spatial attention under simulated shiftwork. Sci Rep. 2020 Sep 11;10(1):14946. doi:10.1038/s41598-020-71800-6 PubMed PMID: 32917940; PubMed Central PMCID: PMC7486912.

36. Aston-Jones G, Rajkowski J, Kubiak P, Valentino RJ, Shipley MT. Role of the locus coeruleus in emotional activation. Prog Brain Res. 1996;107:379–402. doi:10.1016/s0079-6123(08)61877-4 PubMed PMID: 8782532.

37. Posner MI. Measuring alertness. Ann N Y Acad Sci. 2008;1129:193–9. doi:10.1196/annals.1417.011 PubMed PMID: 18591480.

38. Hayat H, Regev N, Matosevich N, Sales A, Paredes-Rodriguez E, Krom AJ, et al. Locus coeruleus norepinephrine activity mediates sensory-evoked awakenings from sleep. Sci Adv. 2020 Apr;6(15):eaaz4232. doi:10.1126/sciadv.aaz4232 PubMed PMID: 32285002; PubMed Central PMCID: PMC7141817.

39. Cowley BR, Snyder AC, Acar K, Williamson RC, Yu BM, Smith MA. Slow Drift of Neural Activity as a Signature of Impulsivity in Macaque Visual and Prefrontal Cortex. Neuron. 2020 Nov 11;108(3):551–567.e8. doi:10.1016/j.neuron.2020.07.021 PubMed PMID: 32810433; PubMed Central PMCID: PMC7822647.

40. Valdez P. Circadian Rhythms in Attention. Yale J Biol Med. 2019 Mar;92(1):81–92. PubMed PMID: 30923475; PubMed Central PMCID: PMC6430172.

41. Aston-Jones G, Rajkowski J, Kubiak P, Valentino RJ, Shipley MT. Role of the locus coeruleus in emotional activation. Prog Brain Res. 1996;107:379–402. doi:10.1016/s0079-6123(08)61877-4 PubMed PMID: 8782532.

42. Samuels ER, Szabadi E. Functional neuroanatomy of the noradrenergic locus coeruleus: its roles in the regulation of arousal and autonomic function part I: principles of functional organisation. Curr Neuropharmacol. 2008 Sep;6(3):235–53. doi:10.2174/157015908785777229 PubMed PMID: 19506723; PubMed Central PMCID: PMC2687936.

43. Gou J, Zhang X, He Y, He K, Xu J. Effects of job demands, job resources, personal resources on night-shift alertness of ICU shift nurses: a cross-sectional survey study based on the job demands-resources model. BMC Nurs. 2024 Sep 12;23(1):648. doi:10.1186/s12912-024-02313-0

44. Vital-Lopez FG, Doty TJ, Reifman J. When to sleep and consume caffeine to boost alertness. SLEEP. 2024 Oct 11;47(10):zsae133. doi:10.1093/sleep/zsae133

45. Plante DT, Hagen EW, Ravelo LA, Peppard PE. Impaired neurobehavioral alertness quantified by the psychomotor vigilance task is associated with depression in the Wisconsin Sleep Cohort study. Sleep Med. 2020 Mar;67:66–70. doi:10.1016/j.sleep.2019.11.1248

46. Joshi S, Gold JI. Pupil Size as a Window on Neural Substrates of Cognition. Trends Cogn Sci. 2020 Jun;24(6):466–80. doi:10.1016/j.tics.2020.03.005 PubMed PMID: 32331857; PubMed Central PMCID: PMC7271902.

47. Joshi S, Li Y, Kalwani RM, Gold JI. Relationships between Pupil Diameter and Neuronal Activity in the Locus Coeruleus, Colliculi, and Cingulate Cortex. Neuron. 2016 Jan 6;89(1):221–34. doi:10.1016/j.neuron.2015.11.028 PubMed PMID: 26711118; PubMed Central PMCID: PMC4707070.

48. Peinkhofer C, Knudsen GM, Moretti R, Kondziella D. Cortical modulation of pupillary function: systematic review. PeerJ. 2019 May 7;7:e6882. doi:10.7717/peerj.6882

49. Wang CA, Baird T, Huang J, Coutinho JD, Brien DC, Munoz DP. Arousal Effects on Pupil Size, Heart Rate, and Skin Conductance in an Emotional Face Task. Front Neurol. 2018;9:1029. doi:10.3389/fneur.2018.01029 PubMed PMID: 30559707; PubMed Central PMCID: PMC6287044.

50. Mitz AR, Chacko RV, Putnam PT, Rudebeck PH, Murray EA. Using pupil size and heart rate to infer affective states during behavioral neurophysiology and neuropsychology experiments. J Neurosci Methods. 2017 Mar 1;279:1–12. doi:10.1016/j.jneumeth.2017.01.004 PubMed PMID: 28089759; PubMed Central PMCID: PMC5346348.

51. Wascher CAF. Heart rate as a measure of emotional arousal in evolutionary biology. Philos Trans R Soc B Biol Sci. 2021 Aug 16;376(1831):20200479. doi:10.1098/rstb.2020.0479

52. Megemont M, McBurney-Lin J, Yang H. Pupil diameter is not an accurate real-time readout of locus coeruleus activity. eLife. 2022 Feb 2;11:e70510. doi:10.7554/eLife.70510

53. Obrist PA, Webb RA, Sutterer JR, Howard JL. CARDIAC DECELERATION AND REACTION TIME: AN EVALUATION OF TWO HYPOTHESES. Psychophysiology. 1970 May;6(6):695–706. doi:10.1111/j.1469-8986.1970.tb02257.x

54. Webb RA, Obrist PA. THE PHYSIOLOGICAL CONCOMITANTS OF REACTION TIME PERFORMANCE AS A FUNCTION OF PREPARATORY INTERVAL AND PREPARATORY INTERVAL SERIES. Psychophysiology. 1970 Jan;6(4):389–403. doi:10.1111/j.1469-8986.1970.tb01749.x

55. Williams DP, Thayer JF, Koenig J. Resting cardiac vagal tone predicts intraindividual reaction time variability during an attention task in a sample of young and healthy adults. Psychophysiology. 2016 Dec;53(12):1843–51. doi:10.1111/psyp.12739 PubMed PMID: 27658566.

56. Schriver BJ, Bagdasarov S, Wang Q. Pupil-linked arousal modulates behavior in rats performing a whisker deflection direction discrimination task. J Neurophysiol. 2018 Oct 1;120(4):1655–70. doi:10.1152/jn.00290.2018

57. De Gee JW, Colizoli O, Kloosterman NA, Knapen T, Nieuwenhuis S, Donner TH. Dynamic modulation of decision biases by brainstem arousal systems. eLife. 2017 Apr 11;6:e23232. doi:10.7554/eLife.23232

58. Nishisato S. Reaction time as a function of arousal and anxiety. Psychon Sci. 1966 Apr;6(4):157–8. doi:10.3758/BF03328005

59. Sassaroli A, Fantini S. Comment on the modified Beer-Lambert law for scattering media. Phys Med Biol. 2004 Jul 21;49(14):N255–257. doi:10.1088/0031-9155/49/14/n07 PubMed PMID: 15357206.

60. Johnston R, Snyder AC, Khanna SB, Issar D, Smith MA. The eyes reflect an internal cognitive state hidden in the population activity of cortical neurons. Cereb Cortex. 2022 Jul 21;32(15):3331–46. doi:10.1093/cercor/bhab418

61. Johnston R, Snyder AC, Schibler RS, Smith MA. EEG Signals Index a Global Signature of Arousal Embedded in Neuronal Population Recordings. eneuro. 2022 May;9(3):ENEURO.0012-22.2022. doi:10.1523/ENEURO.0012-22.2022

62. Warner HR, Russell RO. Effect of combined sympathetic and vagal stimulation on heart rate in the dog. Circ Res. 1969 Apr;24(4):567–73. doi:10.1161/01.res.24.4.567 PubMed PMID: 5780152.

63. Vieira JB, Schellhaas S, Enström E, Olsson A. Help or flight? Increased threat imminence promotes defensive helping in humans. Proc R Soc B Biol Sci. 2020 Aug 26;287(1933):20201473. doi:10.1098/rspb.2020.1473

64. Coventry KR, Constable B. Physiological arousal and sensation-seeking in female fruit machine gamblers. Addiction. 1999 Mar;94(3):425–30. doi:10.1046/j.1360-0443.1999.94342512.x

65. Gordan R, Gwathmey JK, Xie LH. Autonomic and endocrine control of cardiovascular function. World J Cardiol. 2015 Apr 26;7(4):204–14. doi:10.4330/wjc.v7.i4.204 PubMed PMID: 25914789; PubMed Central PMCID: PMC4404375.

66. Grossman P, Taylor EW. Toward understanding respiratory sinus arrhythmia: relations to cardiac vagal tone, evolution and biobehavioral functions. Biol Psychol. 2007 Feb;74(2):263–85. doi:10.1016/j.biopsycho.2005.11.014 PubMed PMID: 17081672.

67. Smith LE. EFFECT OF MUSCULAR STRETCH, TENSION, AND RELAXATION UPON THE REACTION TIME AND SPEED OF MOVEMENT OF A SUPPORTED LIMB. Res Q. 1964 Dec;35:546–53. PubMed PMID: 14276400.

68. Thayer JF, Lane RD. A model of neurovisceral integration in emotion regulation and dysregulation. J Affect Disord. 2000 Dec;61(3):201–16. doi:10.1016/S0165-0327(00)00338-4

69. Kim B, Lee JH, Kang EH, Yu BH. Temperament affects sympathetic nervous function in a normal population. Psychiatry Investig. 2012 Sep;9(3):293–7. doi:10.4306/pi.2012.9.3.293 PubMed PMID: 22993530; PubMed Central PMCID: PMC3440480.

70. Coleman K. Individual differences in temperament and behavioral management practices for nonhuman primates. Appl Anim Behav Sci. 2012 Mar 1;137(3–4):106–13. doi:10.1016/j.applanim.2011.08.002 PubMed PMID: 22518067; PubMed Central PMCID: PMC3327443.

71. Liu Y, Narasimhan S, Schriver BJ, Wang Q. Perceptual Behavior Depends Differently on Pupil-Linked Arousal and Heartbeat Dynamics-Linked Arousal in Rats Performing Tactile Discrimination Tasks. Front Syst Neurosci. 2020;14:614248. doi:10.3389/fnsys.2020.614248 PubMed PMID: 33505252; PubMed Central PMCID: PMC7829454.

72. Liu Y, Rodenkirch C, Moskowitz N, Schriver B, Wang Q. Dynamic Lateralization of Pupil Dilation Evoked by Locus Coeruleus Activation Results from Sympathetic, Not Parasympathetic, Contributions. Cell Rep. 2017 Sep;20(13):3099–112. doi:10.1016/j.celrep.2017.08.094

73. Yang H, Bari BA, Cohen JY, O’Connor DH. Locus coeruleus spiking differently correlates with S1 cortex activity and pupil diameter in a tactile detection task. eLife. 2021 Mar 15;10:e64327. doi:10.7554/eLife.64327

74. Murphy PR, O’Connell RG, O’Sullivan M, Robertson IH, Balsters JH. Pupil diameter covaries with BOLD activity in human locus coeruleus. Hum Brain Mapp. 2014 Aug;35(8):4140–54. doi:10.1002/hbm.22466

75. Wang X, Piñol RA, Byrne P, Mendelowitz D. Optogenetic stimulation of locus ceruleus neurons augments inhibitory transmission to parasympathetic cardiac vagal neurons via activation of brainstem α1 and β1 receptors. J Neurosci Off J Soc Neurosci. 2014 Apr 30;34(18):6182–9. doi:10.1523/JNEUROSCI.5093-13.2014 PubMed PMID: 24790189; PubMed Central PMCID: PMC4004807.

76. Krohn F, Novello M, Van Der Giessen RS, De Zeeuw CI, Pel JJ, Bosman LW. The integrated brain network that controls respiration. eLife. 2023 Mar 8;12:e83654. doi:10.7554/eLife.83654

77. Raut RV, Rosenthal ZP, Wang X, Miao H, Zhang Z, Lee JM, et al. Arousal as a universal embedding for spatiotemporal brain dynamics. Nature. 2025 Sep 24. doi:10.1038/s41586-025-09544-4 PubMed PMID: 40993399.

